# Stress response of membrane-based cell organelles in budding yeast

**DOI:** 10.1101/2024.09.08.611912

**Authors:** Sheng Peng, Bai Li-zhe, Cao Hong, Li Dan

## Abstract

The organelles of yeast demonstrate diverse morphological traits in response to different stress stimuli. However, there is a lack of systematic reports on the structural changes induced by stress stimuli in all membrane-based organelles. Here, we utilized a set of fluorescent protein-based organelle markers to highlight the distinct characteristics of yeast under various stress triggers, including high temperature, hydrogen peroxide, acetic acid, and ethyl alcohol. We found that all of these organelles undergo alterations in structure or function in response to the four stress triggers we tested. Specifically, filamentous mitochondria rupture into smaller segments when exposed to the above four stress conditions. The structure of the endoplasmic reticulum (ER) remains relatively unchanged, but its function is affected. Additionally, high temperature and hydrogen peroxide can induce the Ire1p-mediated ER unfolded protein response (UPR). The translocation of most nuclear-localized proteins to the cytosol is dependent on the specific stress conditions employed. Under the above stress conditions, the vacuole undergoes fusion, resulting in the formation of a larger vacuole from multiple smaller ones. Meanwhile, acetic acid-induced stress leads to the translocation of vacuole-localized proteins Prc1p and Pep4p to unknown puncta, while Ybh3p relocates from the inner vacuole to the vacuole membrane. Proteins localized in the early Golgi, late Golgi, and late endosomes exhibit distinct traits, such as fading away or mis-localization. The structure and function of peroxisomes, lipid droplets, and autophagosomes also undergo modifications. Furthermore, upon exposure to high temperature and ethanol, apoptosis-related proteins Yca1, Aif1, and Mmi1 aggregate instead of remaining dispersed.

## Introduction

Yeast plays a vital role as a cell factory in different industrial biorefineries. However, yeast cells can be harmed by environmental stresses that occur during fermentation, such as high temperatures, ethanol, acidity, and oxidative stress [1]. In response to these extreme conditions, cells undergo programmed cell death, including apoptosis and necrosis. Regardless of the specific molecular mechanisms leading to programmed cell death, apoptotic cells typically display several characteristic features, such as chromatin condensation, nuclear fragmentation, pyknosis, and plasma membrane blebbing [2]. Autophagy can also contribute to cell death under stress, but the exact underlying mechanisms remain unclear [3]. The damage induced by these harmful environmental stresses is regulated by a complex molecular apparatus, involving various components like mitochondria, endoplasmic reticulum, nucleus, lysosomes (vacuoles in yeast), Golgi apparatus and so on [4]. The molecular mechanisms of stress-induced cell response are closely linked to membrane-based organelles.

Mitochondria play a vital role in several key cellular functions, including energy production and programmed cell death (PCD) in eukaryotes. Maintaining a tubular network is crucial for the majority of mitochondrial functions, and this is accomplished through a delicate balance between fusion and fission processes. A notable early feature of apoptosis in both mammalian and yeast cells is the fragmentation of mitochondria into punctuate structures [5, 6]. Apoptosis-inducing factor 1 (AIF1) is a protein located in the mitochondria and is similar to the mammalian AIF. In response to apoptotic insults, Aif1p translocates to the nucleus where it is involved in chromatin condensation and DNA degradation [7]. Nuc1p, which is the yeast counterpart of mammalian endonuclease G (EndoG), is another protein involved in cell death. It also translocates from the mitochondria to the nucleus when apoptosis is induced [8]. NDI1 encodes NADH dehydrogenase, an enzyme found in the inner mitochondrial membrane. Interestingly, Ndi1p is the human equivalent of AMID, which is a protein associated with mitochondria and induces cell death. During Mn-induced apoptosis, Ndi1p can relocate to the cytoplasm [9]. Besides mitochondria, the stress response of the nucleus and ER has also been extensively studied. In yeast, the ER is composed of a network of membranous tubules and sacs known as cisternae. These structures extend throughout the cytoplasm and are connected to the outer membrane of the nuclear envelope, creating a continuous network with the nucleus. During extreme conditions, cells may undergo programmed cell death, with the initial indicators of apoptosis being chromatin condensation and nuclear fragmentation [10]. The ER stress is mediated by the unfolded protein response (UPR), which is characterized by the oligomerization and trans-phosphorylation of inositol-requiring protein-1 (IRE1)[11]. In addition to causing mitochondrial destabilization, nuclear fragmentation and ER stress, lysosomal permeabilization has also been discovered to activate a cell death pathway in mammalian and yeast (vacuole) cells under specific circumstances. Studies in mammalian and yeast systems indicate that the selective permeabilization of lysosomal (vacuole) membranes and the subsequent release of cathepsins or other hydrolases induce apoptosis through a mitochondria-dependent pathway[12–14]. Yeast cathepsin D (CatD), also known as proteinase A (Pep4p), is capable of translocating from the vacuole to the cytosol in response to apoptosis induced by H2O2 or acetic acid [14–16]. Additionally, yeast carboxypeptidase (CPY) Prc1p, also known as proteinase C, along with Pep4p and protein B Prb1p, contributes to the proteolytic function of the vacuole [17]. Mitochondrial outer membrane permeabilization is rigorously controlled by a group of pro- and anti-apoptotic proteins from the BCL-2 family, each of which possesses a BCL-2 homology domain 3 (BH3). Yeast Ybh3p, also known as Bax inhibitor-1 (BXI1), is known to have a BH3 domain and is found in the ER [18] and vacuole [19]. It is capable of relocating from the vacuole to the mitochondria when apoptosis is induced [19].

However, the structural changes of all these important organelles have not been thoroughly investigated under conditions of environmental stress. In this study, our aim was to investigate the structural changes of membrane-based organelles under external stress conditions. Additionally, we wanted to examine the changes in the localization of apoptosis-related proteins in yeast cells when exposed to environmental stress. To achieve this, we utilized multiple fluorescent protein markers for various organelles, including mitochondria, nucleus, ER, Golgi, endosome, peroxisome, and autophagosome. Specifically, we used MitoTracker Red to stain mitochondria and BODIPY (a green lipid dye) to stain lipid droplets. Our findings indicate that exposing yeast cells to high temperature, ethanol, acetic acid, or H2O2 resulted in structural changes in all of these organelles. We used various markers, namely Om14-2GFP, Abf2-2GFP, Afg3-2GFP, Cit1-2GFP, Cox9-mTagBFP, Ndi1-2GFP, Nuc1-2GFP, and MitoTracker, to label the mitochondria structures. Through these markers, we observed that filamentous mitochondria fragmented and transformed into punctate structures. Specifically, we found that two apoptosis-related proteins, Ndi1p and Nuc1p, remained localized to the mitochondria despite the fragmentation. We used Emc1-2GFP, Elo3-mTagBFP, and GFP-HDEL to mark the ER, and found that the signal of the ER resident protein did not change under stress conditions. However, more than half of the signal of GFP-HDEL dispersed from the ER to the cytosol. When examining ER stress using Ire1-GFP, we found that high temperature and oxidative stress could induce Ire1-mediated ER unfolded protein response. The nucleus was marked by Nab2-GFP, Nsr1-GFP, Pus1-2GFP, Nhp6a-2GFP, Nma111-2GFP, Ssn8-2GFP, and Mcd1-2GFP. We found that all of these resident proteins localized in the nucleus under normal conditions, but relocated to the cytosol partially or completely under external stresses. The vacuole was marked by Pho8-GFP, Ybh3-2GFP, Prc1-2GFP, and Pep4-2GFP. Under normal conditions, all of these vacuole markers were localized within the vacuole, with the vacuole membrane signal of Pho8 being higher than the others. However, under acetic acid-induced conditions, the GFP signal of Ybh3p moved from the inner vacuole to the vacuole membrane, and the GFP signal of Prc1p and Pep4 transferred from the inner vacuole to several unknown puncta. High temperature, ethanol, and oxidative stress did not change the localization of any of the four vacuole markers we tested. We also examined the early-Golgi, late-Golgi, late-endosome, lipid droplet, peroxisome, and autophagosome using their resident proteins or by staining with dye. In addition to these organelles, we also re-examined the localization of three apoptosis-relevant proteins, Yca1p, Aif1p, and Mmi1p.

## Materials and Methods

### Yeast strains, media and culture conditions

All strains used in this study are based on the *S. cerevisiae* strain BY4741 (*MATa his3Δ1 leu2Δ0 ura3Δ0 met15Δ0*). The strains used and created in this study are summarized in Table 1. Yeast cells were grown in complex medium (YPD) containing 1% (w/v) yeast extract, 2% (w/v) peptone, and 2% (w/v) D-glucose. Solid media were prepared by adding 2% (w/v) agar. SD-N medium was used for nitrogen starvation, consisting of 0.17% (w/v) yeast nitrogen base without amino acids and ammonium sulfate, and 2% (w/v) D-glucose. SMD medium (2% glucose, 0.67% yeast nitrogen base without amino acids, 30 mg/liter adenine, 30 mg/liter lysine, 30 mg/liter methionine, 20 mg/liter histidine, 20 mg/liter uracil, 50 mg/liter tryptophan, 50 mg/liter leucine) was used to wash and resuspend cells before the fluorescent microscopy experiment.

**Table 1.** Strains used in this study.

| Name | Genotype |
| --- | --- |
| BY4741 | *MATa his3Δ1 leu2Δ0 met15Δ0 ura3Δ0* |
| YB1 | *BY4741 OM14::OM14-2GFP-URA3 (K.l.)* |
| YB2 | *BY4741 CIT1::CIT1-2GFP-URA3(K.l.)* |
| YB3 | *BY4741 AFG3::AFG3-2GFP-URA3(K.l.)* |
| YB4 | *BY4741 ABF2::ABF2-2GFP-URA3(K.l.)* |
| YB5 | *BY4741 COX9::COX9-mTagBFP-kanMX NDI1::NDI1-2GFP-URA3(K.l.)* |
| YB6 | *BY4741 COX9::COX9-mTagBFP-kanMX NUC1::NUC1-2GFP-URA3(K.l.)* |
| YB7 | *BY4741 ELO3::ELO3-mTagBFP-kanMX EMC1::EMC1-2GFP-URA3(K.l.)* |
| YB8 | *BY4741 pTPI1:: pTPI1-SS-GFP-HDEL-URA3 (K.l.)* |
| YB9 | *BY4741 IRE1::IRE1-2GFP-URA3(K.l.)* |
| YB10 | *BY4741 ELO3::ELO3-mTagBFP-kanMX NAB2::NAB2-GFP-URA3(K.l.)* |
| YB11 | *BY4741 NSR1::NSR1-GFP-URA3(K.l.)* |
| YB12 | *BY4741 PUS1::PUS1-2GFP-URA3(K.l.)* |
| YB13 | *BY4741 NHP6A::NHP6A-2GFP-URA3(K.l.)* |
| YB14 | *BY4741 SSN8::SSN8-2GFP-URA3(K.l.)* |
| YB15 | *BY4741 NMA111::NMA111-2GFP-URA3(K.l.)* |
| YB16 | *BY4741 MCD1::MCD1-2GFP-URA3(K.l.)* |
| YB17 | *BY4741 PHO8::pCU-GFP-PHO8-URA3(K.l.)* |
| YB18 | *BY4741 YBH3::YBH3-2GFP-URA3(K.l.)* |
| YB19 | *BY4741 PRC1::PRC1-2GFP-URA3(K.l.)* |
| YB20 | *BY4741 PEP4::PEP4-2GFP-URA3(K.l.)* |
| YB21 | *BY4741 CHS5::CHS5-GFP-URA3(K.l.)* |
| YB22 | *BY4741 SEC7::SEC7-2GFP-URA3(K.l.)* |
| YB23 | *BY4741 VRG4::VRG4-2GFP-URA3(K.l.)* |
| YB24 | *BY4741 SYS1::SYS1-2GFP-URA3(K.l.)* |
| YB25 | *BY4741 SNF7::SNF7-GFP-URA3(K.l.)* |
| YB26 | *BY4741 VPS4::VPS4-GFP-URA3(K.l.)* |
| YB27 | *BY4741 PEX1::PEX1-2GFP-URA3(K.l.)* |
| YB28 | *BY4741 ATG8::pATG8-GFP-ATG8-URA3(K.l.)* |
| YB29 | *BY4741 COX9::COX9-mTagBFP-kanMX YCA1::YCA1-2GFP-URA3(K.l.)* |
| YB30 | *BY4741 COX9::COX9-mTagBFP-kanMX AIF1::AIF1-2GFP-URA3(K.l.)* |
| YB31 | *BY4741 ELO3::ELO3-mTagBFP-kanMX MMI1::pMMI1-2GFP-MMI1-URA3(K.l.)* |

For high temperature experiments, yeast cells were grown in YPD medium until reaching an OD 600 of 0.7 at 30℃, and then shifted to 42℃. In acetic acid-induced experiments, yeast cells were grown in YPD medium until reaching an OD 600 of 0.7 at 30℃, and acetic acid was added to the logarithmic phase medium at various final concentrations (0.1% (16.7 mM), 0.2%, 0.4%, and 0.6% (v/v)). Similarly, in ethanol-induced experiments, yeast cells were grown in YPD medium until reaching an OD 600 of 0.7 at 30℃, and ethanol was added to the logarithmic phase medium at different final concentrations (2%, 4%, 8%, and 10% (v/v)). For H2O2-induced experiments, yeast cells were grown in YPD medium until reaching an OD 600 of 0.7 at 30℃, and H2O2 was added to the logarithmic phase medium at various final concentrations (0.5, 1.0, 2.0, and 4.0 mM).

### Plasmid and Cloning experiments

Some of the plasmids used to tag cell organelle markers were created in our previous study [20]. The remaining plasmids were constructed using standard methods. This involved amplifying fragments through PCR, digesting them with restriction enzymes, and subsequently joining them together. Prior to transforming them into cells, linear fragment were obtained using restriction enzymes. The majority of the plasmids were designed as 3’-knock-in constructs. Only GFP-Pho8, GFP-Atg8 and 2GFP-Mmi1 were designed as an additional copy. The plasmids used in this work are listed in Table 2.

**Table 2.** Plasmids constructed in this study.

| **Plasmid** | **Parental plasmid** | **Genome integration locus** |
| --- | --- | --- |
| OM14-2GFP-ter-URA3 | UG72-2GFP | C terminal |
| CIT1-2GFP-ter-URA3 | UG72-2GFP | C terminal |
| AFG3-2GFP-ter-URA3 | UG72-2GFP | C terminal |
| ABF2-2GFP-ter-URA3 | UG72-2GFP | C terminal |
| COX9-mTagBFP-ter-kanMX | UG6- mTagBFP | C terminal |
| NDI1-2GFP-ter-URA3 | UG72-2GFP | C terminal |
| NUC1-2GFP-ter-URA3 | UG72-2GFP | C terminal |
| ELO3-mTagBFP-ter-kanMX | UG6- mTagBFP | C terminal |
| EMC1-2GFP-ter-URA3 | UG72-2GFP | C terminal |
| pTPI1-SS-GFP-HDEL-URA3 | UG72-GFP | TPI1 promoter |
| IRE1-2GFP-ter-URA3 | UG72-2GFP | C terminal |
| NAB2-GFP-ter-URA3 | UG72-GFP | C terminal |
| NSR1-GFP-ter-URA3 | UG72-GFP | C terminal |
| PUS1-2GFP-ter-URA3 | UG72-2GFP | C terminal |
| NHP6A-2GFP-ter-URA3 | UG72-2GFP | C terminal |
| SSN8-2GFP-ter-URA3 | UG72-2GFP | C terminal |
| NMA111-2GFP-ter-URA3 | UG72-2GFP | C terminal |
| MCD1-2GFP-ter-URA3 | UG72-2GFP | C terminal |
| pCU-GFP-PHO8-ter-URA3 | UG72-2GFP | PHO8 ORF |
| YBH3-2GFP-ter-URA3 | UG72-2GFP | C terminal |
| PRC1-2GFP-ter-URA3 | UG72-2GFP | C terminal |
| PEP4-2GFP-ter-URA3 | UG72-2GFP | C terminal |
| CHS5-GFP-ter-URA3 | UG72-GFP | C terminal |
| SEC7-2GFP-ter-URA3 | UG72-2GFP | C terminal |
| VRG4-2GFP-ter-URA3 | UG72-2GFP | C terminal |
| SYS1-2GFP-ter-URA3 | UG72-2GFP | C terminal |
| SNF7-GFP-ter-URA3 | UG72-GFP | C terminal |
| VPS4-GFP-ter-URA3 | UG72-GFP | C terminal |
| PEX1-2GFP-ter-URA3 | UG72-2GFP | C terminal |
| pATG8-GFP-ATG8-ter-URA3 | UG72-2GFP | ATG8 promoter |
| YCA1-2GFP-ter-URA3 | UG72-2GFP | C terminal |
| AIF1-2GFP-ter-URA3 | UG72-2GFP | C terminal |
| pMMI1-2GFP-MMI1-ter-URA3 | UG72-2GFP | C terminal |

### Fluorescence microscopy

1 ml of liquid culture per sample was washed and resuspended in SMD medium. The resuspended sample was then placed on concanavalin A-coated glass bottom 35mm petri dishes and allowed to precipitate for 2 minutes. Then, 1 ml of New SMD medium was added to the dishes, and image stacks (15 slices, 0.5μm step size) were collected using an Olympus IX83 inverted fluorescence microscope equipped with a Hamamatsu Orca Flash4.0 LT camera and a Lumencor Spectra X six-channel light source. The excitation intensity was set to 100%, and the exposure time for each frame was 100 ms for GFP- and BFP-tagged proteins.

For BODIPY (lipid droplet) and MitoTracker red (mitochondria) staining, 1 μl of dye (0.01 mg/ml DAPI or 0.1mM MitoTracker red) was added to 100μl of yeast liquid culture, and the mixture was left to sediment on concanavalin A-coated cover dishes for 2 min. After washing with fresh SMD medium and re-adding new SMD, adhered yeast cells were observed under the microscope. The excitation intensity was set to 100%, and the exposure time for BODIPY and MitoTracker red staining was 100 ms.

## Results and discussion

### Mitochondrion

Mitochondria are highly dynamic, constantly undergoing fission and fusion, which helps maintain their function and integrity. To systematically study the effect of environmental stress on the structure of membrane-based organelles in yeast, we first labeled the mitochondria using fluorescent protein-based mitochondrial proteins. Om14p is a key protein in the outer mitochondrial membrane of yeast cells, primarily involved in the import of mitochondrial proteins [21]. Abf2p is a mitochondrial DNA binding protein involved in mitochondrial DNA replication and recombination [22]. Afg3p is a component of the mitochondrial inner membrane m-AAA protease [23]. Cit1p, the mitochondrial citrate synthase, plays a crucial role in the citric acid cycle by catalyzing the conversion of acetyl-CoA and oxaloacetate into citrate [24]. Cox9p (Cytochrome c oxidase subunit 9) is a component of the cytochrome c oxidase (Complex IV) of the mitochondrial respiratory system [25]. Ndi1p is a type of NADH dehydrogenase that catalyzes the transfer of electrons from NADH to ubiquinone (coenzyme Q) in the mitochondrial electron transport chain [26]. Nuc1p is an endo/exonuclease enzyme that is localized in the mitochondrial matrix and functions to degrade damaged or unneeded mitochondrial DNA [27].

To more accurately observe changes in mitochondrial structure under stress conditions, we selected several mitochondrial proteins and dyes for labeling and observation. Among all the candidate constructs, we observed that Om14-2GFP, Abf2-2GFP, Afg3-2GFP, Cit1-2GFP, Cox9-mTagBFP, Ndi1-2GFP, Nuc1-2GFP and MitoTracker displayed a tubular mitochondrial network during the log phase at 30℃ in YPD medium (Figure 1 A-J). Upon shifting the yeast cells from 30℃ to 42℃ for more than 5 minutes, the tubular mitochondria began to fragment (Figure 1 A). After twenty minutes, they had completely fragmented (Figure 1 A). Besides high temperature, the fragmentation of mitochondria can also be caused by the metabolic byproduct of acetic acid and ethanol, as well as oxidative stress. Our findings indicate that when the concentration of acetic acid exceeds 0.2% and ethanol exceeds 10% in the medium, the filamentous mitochondria of yeast cells are completely disrupted (Figure 1 B-C). We also observed a similar phenomenon under H_2_O_2_-induced conditions (Figure 1 D). To accurately observe the behavior of mitochondria, we used MitoTracker, a specific dye that stains mitochondria. This dye helps highlight the actual structure of mitochondria under different induction conditions (Figure 1 H). Our findings indicate that both the MitoTracker and all the GFP or mTagBFP labeled mitochondrial proteins exhibited similar structural changes when exposed to high temperature, acetic acid, ethanol, and H2O2-induced conditions (Figure 1 A-J). Previous studies have shown that the mitochondria NADH dehydrogenase Ndi1p can relocate to the cytoplasm during Mn-induced apoptosis [9]. However, in this study, we consistently observed that Ndi1-2GFP remains within the mitochondria under high temperature, acetic acid, ethanol, and H2O2-induced conditions (Figure 1 I). It has been reported that Nuc1p, the yeast EndoG, becomes concentrated in the nuclei after treatment with 0.4 mM H2O2 [8]. Nonetheless, we observed complete colocalization of Nuc1-2GFP and Cox9-mTagBFP under conditions induced by high temperature, acetic acid, ethanol, and H2O2 (Figure1 J).

**Fig 1.**
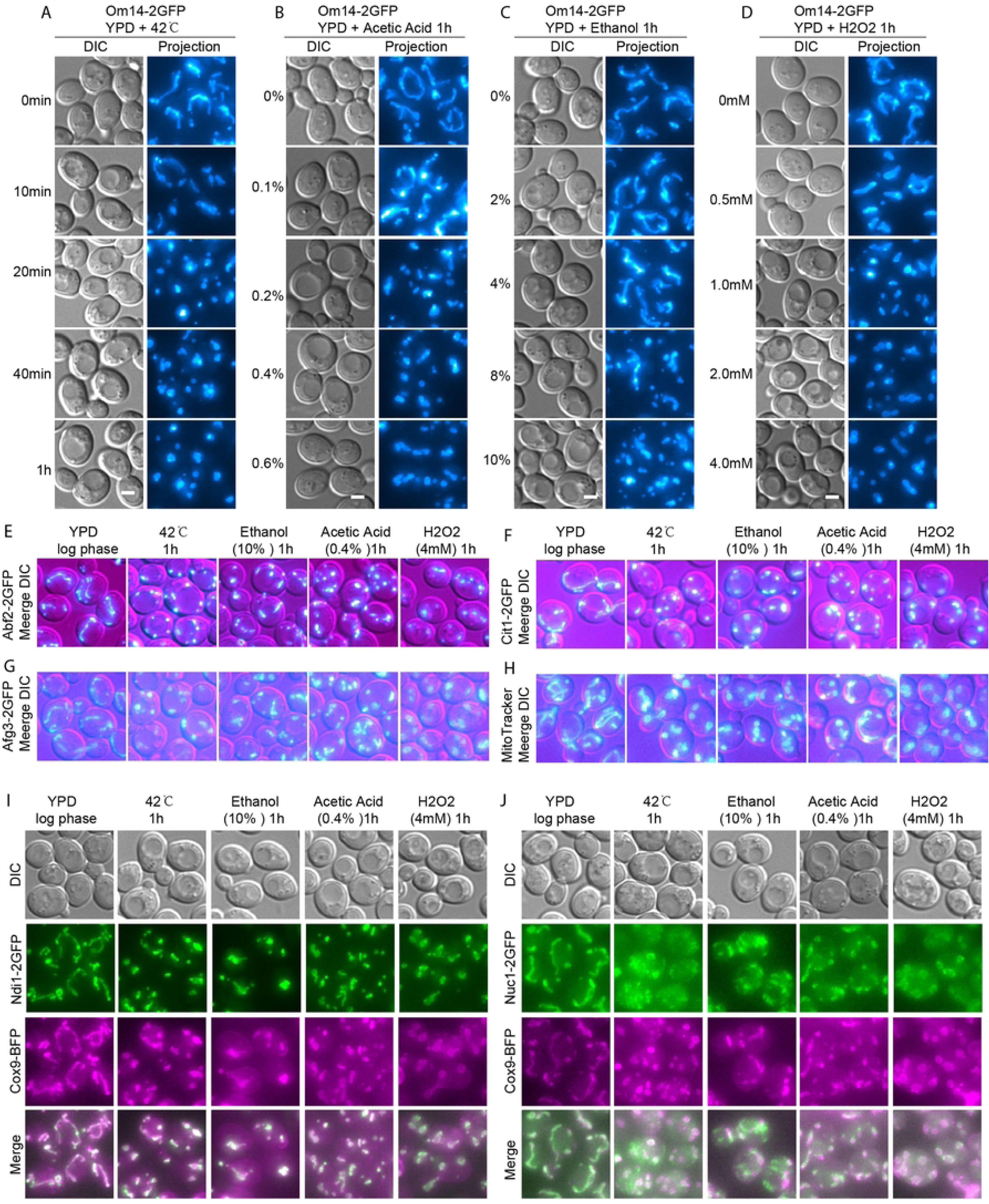
Stress response of mitochondria. Mil-log-phase yeast cells expressing the indicated fluorescent protein fusion constructs were induced by high temperature, acetic acid, ethanol, and H_2_O_2_, and then immobilized on concanavalin A-coated petri dishes. Z-stacks were captured, consisting of 15 slices with a step size of 0.5 μm. For colocalization experiments, the images shown are slices. Projection refers to the maximal intensity projection of z-stacks, while DIC stands for differential interference contrast. (A-D) Structural changes of mitochondria (Om14-2GFP) induced by high temperature, acetic acid, ethanol, and H_2_O_2_. (E-H) Structural changes induced by high temperature, acetic acid, ethanol, and H_2_O_2_ were observed in DNA binding protein Abf2 (E), citrate synthase Cit1p (F), mitochondrial inner proteins Afg3 (G), and red mitochondria marker (H). (I) Colocalization between mitochondrial protein Cox9p and Ndi1p. (J) Colocalization between mitochondrial protein Cox9p and Nuc1p

### Endoplasmic Reticulum (ER)

The endoplasmic reticulum (ER) is a vital organelle in eukaryotic cells that performs essential functions in the synthesis, folding, modification, and transport of proteins and lipids. In yeast cells, the ER can be divided into three distinct parts when viewed under a light microscope: the nuclear ER, which is continuous with the nuclear envelope surrounding the nucleus; the cortical ER, located beneath the plasma membrane and closely associated with the cell’s periphery; and the intermediate ER structures that connect the nuclear ER and the cortical ER, forming a network of tubules and sheets throughout the cytoplasm. EMC1 is a core component of the Endoplasmic Reticulum Membrane Protein Complex (EMC), which plays a crucial role in the biogenesis and proper folding of transmembrane proteins within the ER[28]. ELO3 is an enzyme involved in the elongation of saturated and unsaturated very long-chain fatty acids (VLCFAs) in the ER[29]. HDEL is a retention signal found at the C-terminus of certain proteins in the ER [30]. The Emc1-2GFP, Elo3-mTagBFP, and GFP-HDEL all labeled the three parts of the ER (Figure 2 A). Our results from live-cell imaging showed that the ER structure labeled by Emc1-2GFP and Elo3-mTagBFP remained relatively stable and colocalized with each other during exposure to high temperature, acetic acid, ethanol, and H2O2 (Figure 2 A). However, the signal labeled with GFP-HDEL disappeared significantly, especially under high temperature conditions (Figure 2 B).

**Fig 2.**
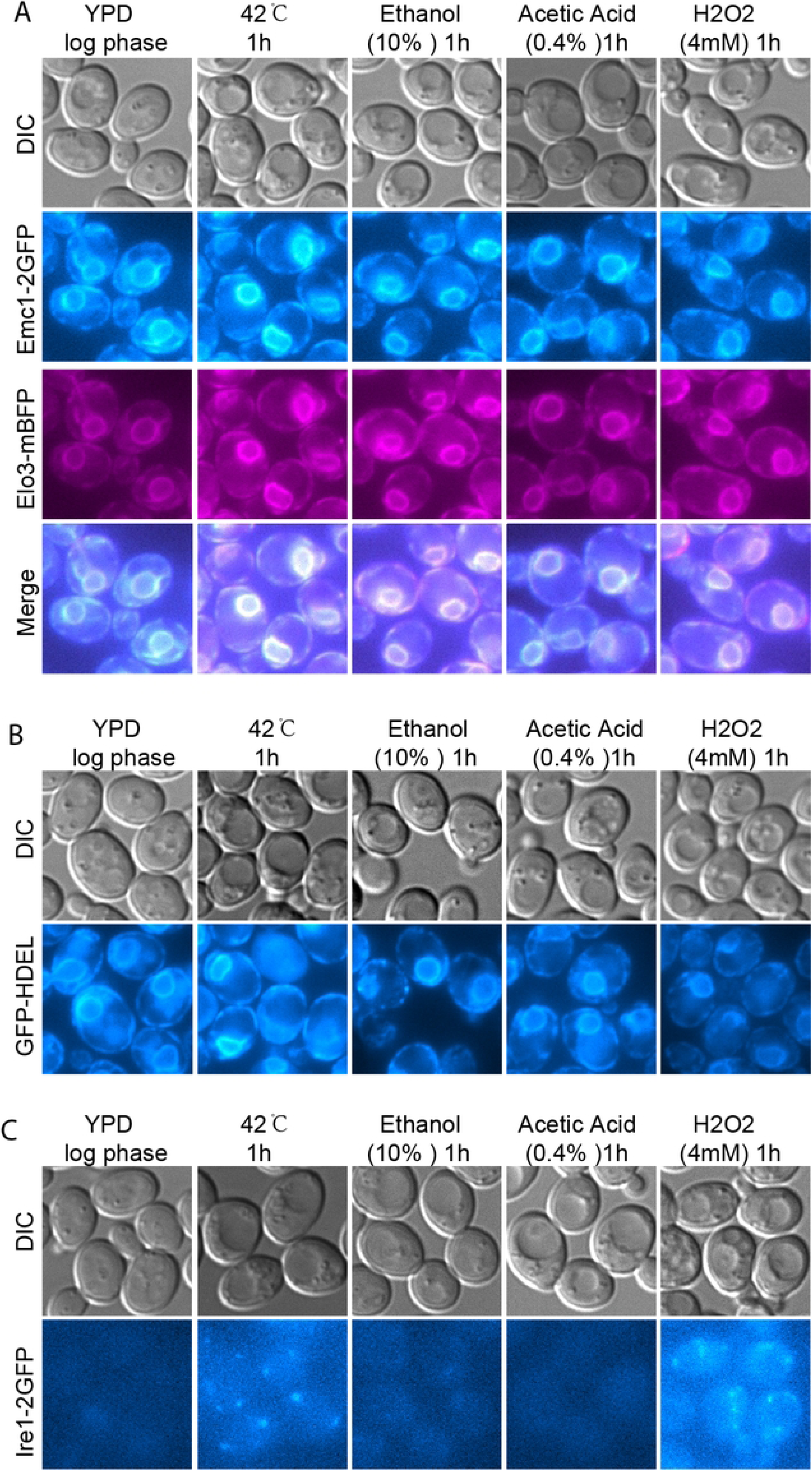
Stress response of ER. Images were captured and presented as in Fig 1. (A-B) The structure of the ER was indicated by Emc1-2GFP, Elo3-mTagBFP and GFP-HDEL under high temperature, acetic acid, ethanol, and H2O2-induced conditions. Colocalization between the green and red ER markers (A). (C) The unfolded protein response of ER was observed by Ire1-2GFP.

The Unfolded Protein Response (UPR) is a cellular stress response pathway that is activated when unfolded or misfolded proteins accumulate in the endoplasmic reticulum (ER). The UPR is crucial for maintaining cellular homeostasis, especially in conditions that disrupt protein folding in the ER. In yeast, IRE1 plays a key role in the ER stress response as the primary sensor and activator of the UPR. [31]. When there is an accumulation of misfolded proteins in the ER, IRE1 undergoes oligomerization and autophosphorylation, which activates its RNase domain [32]. In this work, we found that only high temperature and H2O2 can induce the oligomerization of Ire1p (Figure2 C).

### Nucleus

The yeast nucleus is surrounded by a double membrane called the nuclear envelope, which separates the nucleus from the cytoplasm. The nuclear envelope is punctuated with nuclear pores that regulate the transport of molecules in and out of the nucleus. Nab2p (Nuclear poly(A) binding protein 2) is an essential nucleus protein in yeast that plays a critical role in mRNA metabolism, particularly in the processing, export, and stability of messenger RNA (mRNA)[33]. Nsr1p (Nucleolar Seroine-rich protein 1) is a nucleolar protein in yeast that plays a crucial role in ribosome biogenesis, particularly in the processing and assembly of ribosomal RNA (rRNA)[34]. Pus1p (Pseudouridine Synthase 1) is an enzyme in yeast that is involved in the post-transcriptional modification of RNA[35]. Nhp6ap is a high-mobility group (HMG) protein in yeast, and it plays a crucial role in chromatin structure and function[36]. Nma111p is a nuclear serine protease that is primarily localized in the nucleus, where it exerts its protease activity on nuclear substrates involved in the apoptotic pathway[37]. Ssn8/Srb11 is a component of the CDK8 module within the Mediator complex in nucleus. It plays a crucial role in the negative regulation of transcription, particularly in response to stress and during certain stages of the cell cycle[38]. Mcd1, also known as Scc1 (Sister chromatid cohesion 1), is a crucial protein in nucleus involved in the process of sister chromatid cohesion during cell division[39].

Among the seven candidate constructs for the nucleus, we observed that Nab2-GFP, Nsr1-GFP, Pus1-GFP, Nhp6a-2GFP, Nma111-2GFP, Ssn8-2GFP, and Mcd1-2GFP localized to the nucleus under log phase in YPD medium (Figure 3 A-D). However, when exposed to high temperatures at 42℃, the GFP signal of Nab2p, Pus1p, Nma111p, and Mcd1p decreased by more than half (Figure 3 A-D). Under ethanol-induced conditions, the nuclear GFP signal of Nab2p decreased significantly, while the other six proteins showed only a slight decrease (Figure 3 A-D). No changes were observed in the nucleus signal of any of the seven proteins under acetic acid-induced conditions (Figure 3 A-D). Finally, under H2O2-induced conditions, the GFP signal of Nab2p disappeared completely into the cytoplasm, and the GFP signal of the other six candidate proteins also weakened (Figure 3 A-D).

**Fig 3.**
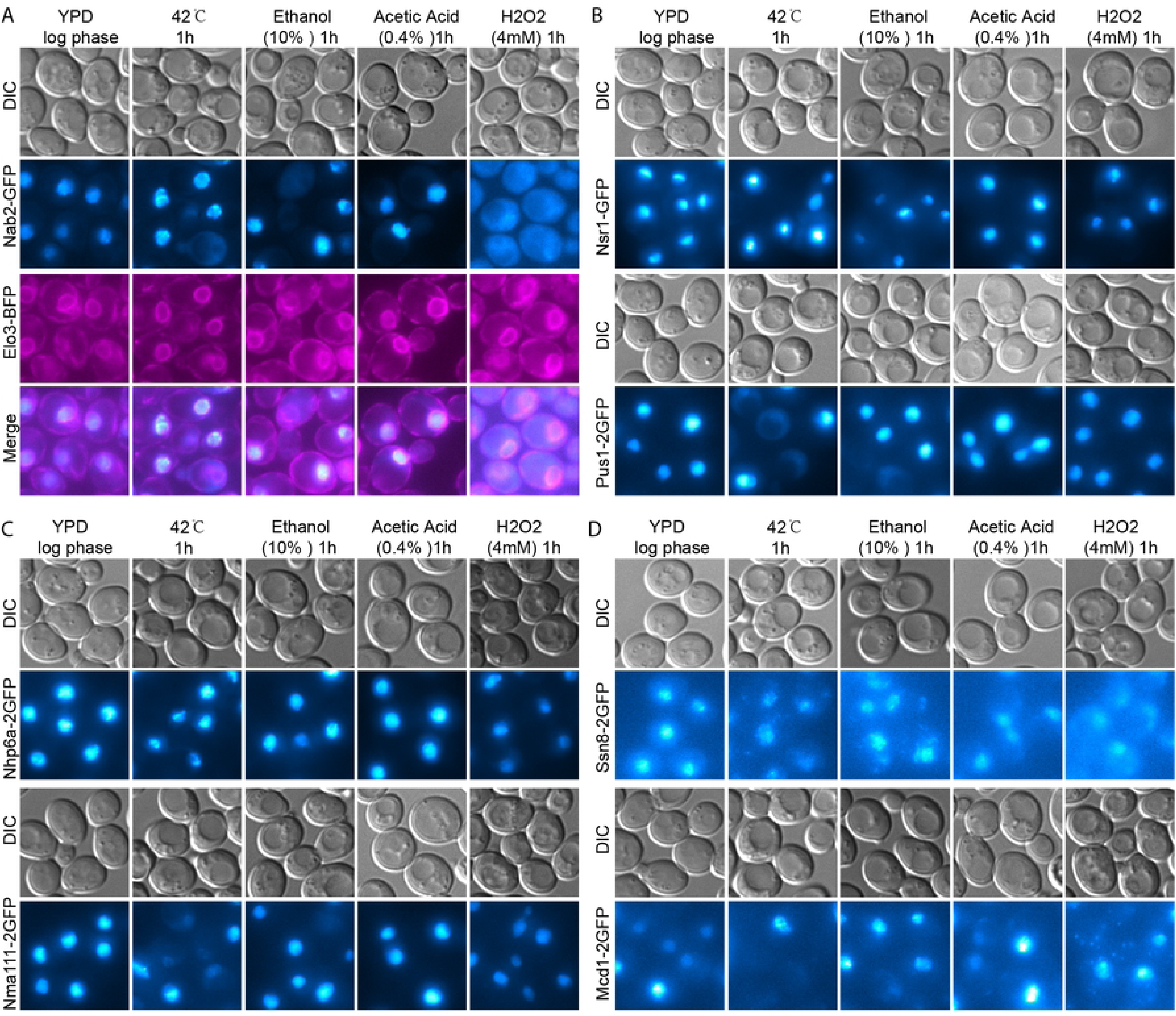
Stress response of nucleus. Images were captured and presented as in Fig 1. (A-D) The structure of the nucleus was indicated by Nab2-2GFP, Nsr1-GFP, Pus1-2GFP, Nhp6a-2GFP, Nma111-2GFP, Ssn8-2GFP, and Mcd1-2GFP under high temperature, acetic acid, ethanol, and H2O2-induced conditions. Colocalization between the green and red markers (A).

### Vacuoles

In yeast cells, vacuoles are organelles similar to lysosomes in animal cells. They have multiple functions, including storing ions like calcium and phosphate, amino acids, and other nutrients. Vacuoles in yeast also play a crucial role in protein degradation and the cell’s response to environmental stresses such as changes in nutrient availability or osmotic pressure. One specific enzyme called Pho8p, which is a vacuolar (lysosomal) alkaline phosphatase, is essential for phosphate metabolism and regulation[40]. Another protein called BXI1, also known as Ybh3p in yeast, contains a BCL-2 homology (BH3) domain. Previous work showed that treatment with acetic acid, BXI1/Ybh3 translocate from predominantly vacuolar sites to mitochondria[19] Prc1p, also known as vacuolar carboxypeptidase Y, is a vital enzyme in yeast that degrades proteins and peptides within the vacuole[41]. Another enzyme called Pep4p, which is encoded by the PEP4 gene in yeast, is responsible for protein degradation and activating other vacuolar hydrolases. Pep4p has a significant role in autophagy, stress adaptation, and overall protein turnover within the cell [42]. Among the four candidate constructs for the vacuoles, we observed that GFP-Pho8, Ybh3-2GFP, Prc1-2GFP, and Pep4-2GFP showed localization to the vacuole under log phase conditions (Figure 4 A-D). The GFP-Pho8 signal did not under environmental stress conditions such as high temperature, acetic acid, and H2O2 (Figure 4 A). However, the Ybh3-2GFP predominantly translocated to the vacuolar membrane under acetic acid induced conditions (Figure 4 B). On the other hand, the vacuolar localization of Prc1-2GFP and Pep4-2GFP completely disappeared, and only some puncta were observed in the cytoplasm under acetic acid induced conditions (Figure 4 C-D). The localization of Ybh3-2GFP, Prc1-2GFP, and Pep4-2GFP did not change much under high temperature, ethanol, and H2O2 conditions and exhibited a fusion pattern similar to GFP-Pho8 (Figure 4 A-D).

**Fig 4.**
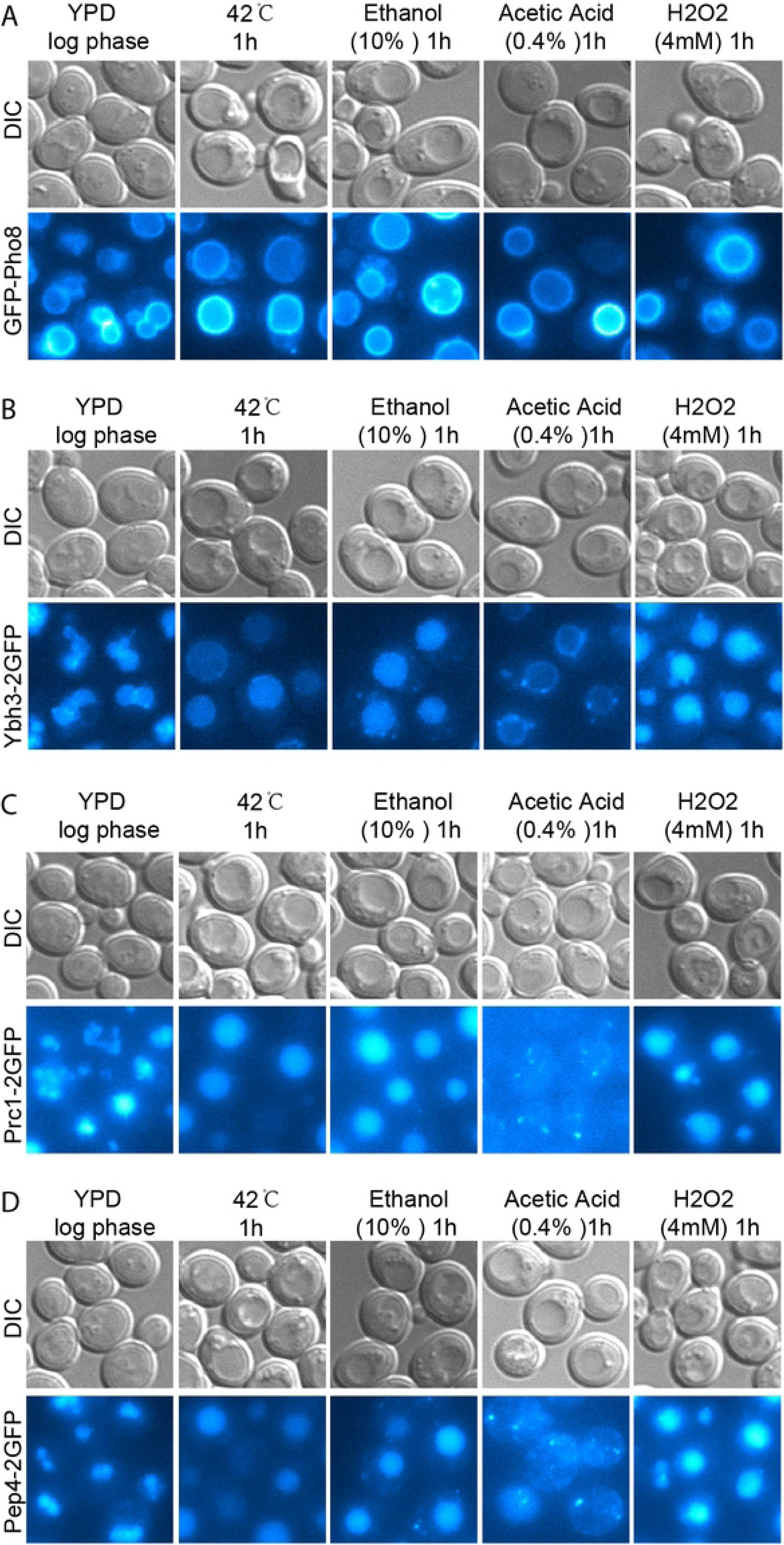
Stress response of vacuole. Images were captured and presented as in Fig 1. (A-D) The structure of the nucleus was indicated by Nab2-2GFP, Nsr1-GFP, Pus1-2GFP, Nhp6a-2GFP, Nma111-2GFP, Ssn8-2GFP, and Mcd1-2GFP under high temperature, acetic acid, ethanol, and H2O2-induced conditions. Colocalization between the green and red markers (A).

### Golgi and endosome

The Golgi apparatus is divided into different functional regions. The early Golgi, specifically the cis-Golgi network (CGN), receives and initiates the processing of cargo from the ER. The late Golgi, mainly the trans-Golgi network (TGN), is responsible for the final sorting and packaging of cargo into specific vesicles. Its function is to ensure that proteins and lipids are accurately delivered to their respective destinations within the cell. Endosomes, which are membrane-bound vesicles, are part of the endocytic pathway. They manage the internalization and sorting of various materials, including those from the Golgi. Vrg4p is a GDP-mannose transporter found on the early Golgi membrane [43]. Sys1p is an early Golgi-associated protein in yeast that plays a crucial role in vesicle trafficking and the localization of Golgi enzymes [44]. Chs5p is an essential protein in Saccharomyces cerevisiae that is responsible for the late-Golgi-to-plasma membrane trafficking of specific cargo proteins, particularly those involved in chitin synthesis[45]. Sec7p, localized in the late Golgi, is a central player in the secretory pathway. It facilitates the formation of vesicles that transport cargo proteins to their appropriate destinations within the cell [46]. Snf7p and Vps4p are two proteins in Saccharomyces cerevisiae that are key components of the Endosomal Sorting Complex Required for Transport (ESCRT) machinery. Both of these proteins are found in late endosomes [20].

Among the candidate constructs, we observed that the GFP signal of the early Golgi proteins Vrg4-2GFP and Sys1-2GFP diminished significantly during high temperature and ethanol-induced conditions (Figure 5 A-B). Additionally, under acetic acid and H2O2 conditions, the number of GFP puncta for both Vrg4 and Sys1 decreased (Figure 5 A-B). Moving on to the late Golgi, we discovered that the GFP signal of Chs5 and Sec7 completely vanished under H2O2-induced conditions. However, under high temperature conditions, the GFP puncta clustered together, and under ethanol-induced conditions, the GFP signal faded significantly. Moreover, under acetic acid-induced conditions, the GFP puncta became larger and lighter, especially in the case of Sec7-2GFP (Figure 5 C-D). The late endosome candidates Snf7-GFP and Vps4-GFP exhibited distinct signals in this study. Specifically, we observed that Snf7-GFP moved from aggregated puncta to the vacuole membrane during high temperature, ethanol, acetic acid, and H2O2 conditions (Figure 5 E). Furthermore, the GFP signal of Vps4-GFP significantly disappeared during high temperature, ethanol, and acetic acid-induced conditions, and to a lesser extent during H2O2 conditions (Figure 5 F).

**Fig 5.**
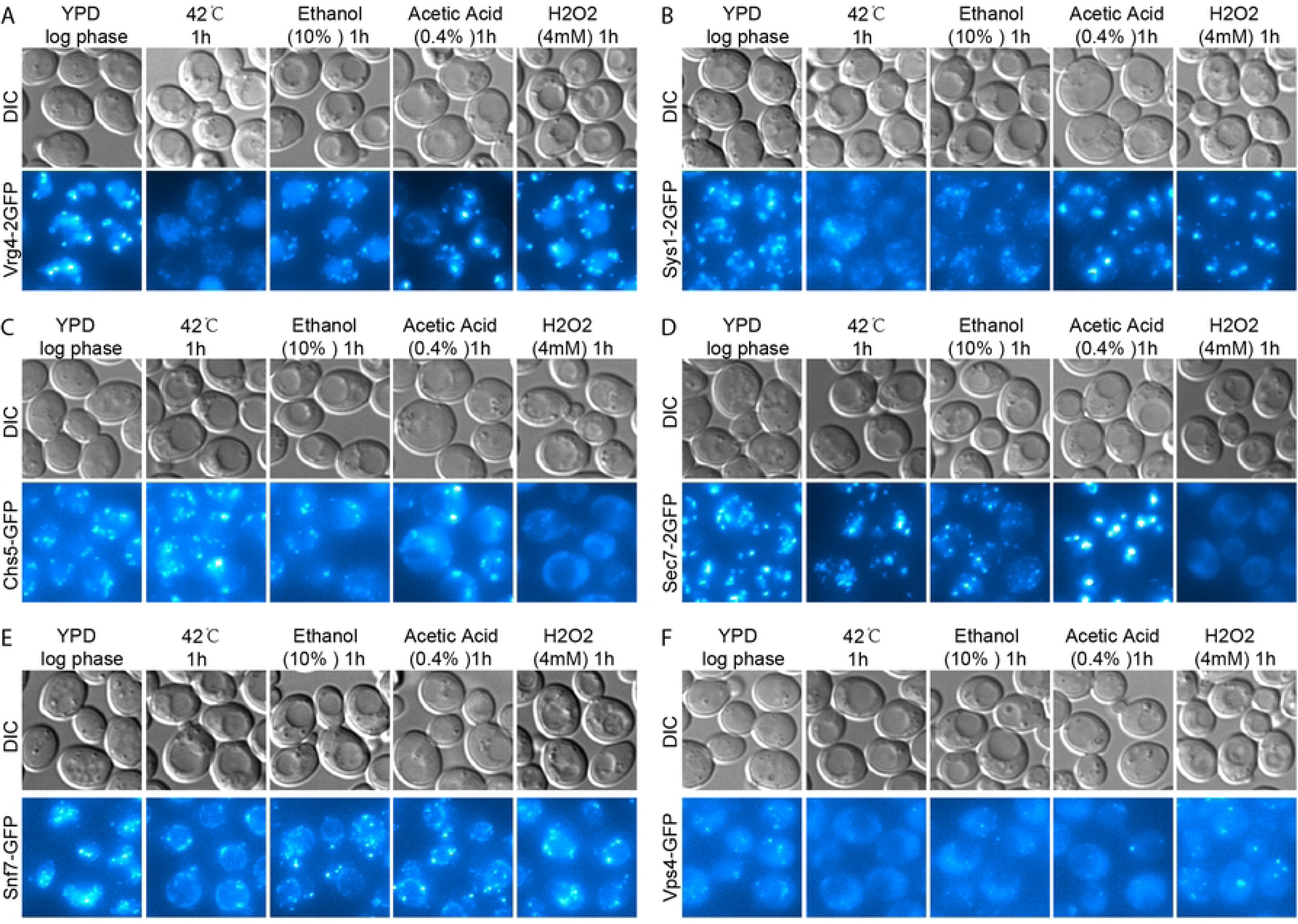
Stress response of Golgi and endosomes. Images were captured and presented as in Fig 1 (A-F). The structure of the early-Golgi was indicated by Vrg4-2GFP and Sys1-2GFP (A-B), While the structure of the late-Golgi was indicated by Chs5-GFP and Sec7-2GFP (C-D). Additionally, the structure of the late-endosome was indicated by Chs5-GFP and Sec7-2GFP (E-F).

### Lipid droplet, peroxisome, and autophagosome

Lipid droplets play a crucial role in lipid metabolism, energy storage, and lipid homeostasis. They are surrounded by a monolayer of phospholipids and proteins, which distinguishes them from other organelles that typically have a bilayer membrane. Abnormal accumulation of lipid droplets is associated with various diseases, including obesity, diabetes, and liver diseases. BODIPY dyes are non-toxic and can be used to stain lipid droplets in live cells [47]. In this study, we observed a significant decrease in lipid droplets under high temperature conditions (Figure 6A). Additionally, acetic acid and H2O2 could induce the aggregation of lipid droplet (Figure 6A). Ethanol had only a minor effect on lipid droplets (Figure 6A).

**Fig 6.**
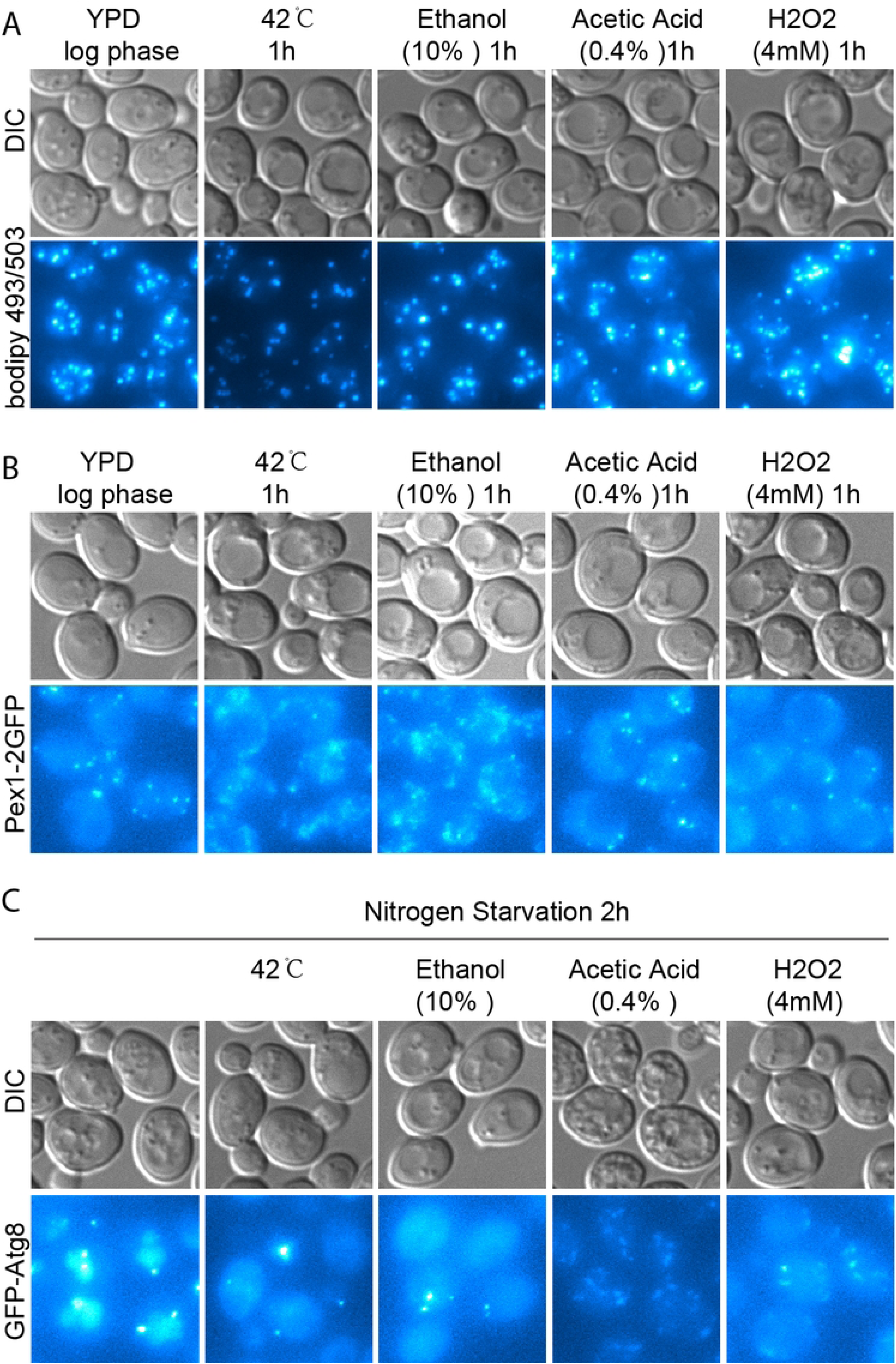
Stress response of lipid droplet, peroxisome, and autophagosome. Images were captured and presented as in Fig 1 (A-C). The structure of lipid droplets was indicated by green dye BODIPY (A), While the structure of the peroxisome was indicated by Pex1-2GFP (B). Additionally, the structure of the autophagosome was indicated by GFP-Atg8 (E-F).

Peroxisomes are small organelles surrounded by a single lipid bilayer membrane. They contain enzymes that are vital for their functions. Unlike lipid droplets, peroxisomes have a specific structure. They play a significant role in various metabolic processes, particularly in breaking down very-long-chain fatty acids through beta-oxidation. Pex1p is an ATPase that is crucial for importing peroxisomal matrix proteins, which are essential for peroxisome function and maintenance[48]. In our study, we observed that the Pex1-2GFP puncta remained relatively unchanged under acetic acid and H2O2 conditions (Figure 6 B). However, both high temperature and ethanol induced the aggregation of Pex1-2GFP puncta (Figure 6 B).

Autophagy is a cellular process that involves breaking down and recycling unnecessary or damaged parts of cells. It plays a crucial role in maintaining cell balance, responding to stress, and ensuring the quality of cellular components. Atg8p, the main player in autophagy, binds to phosphatidylethanolamine (PE) on the autophagosome membrane, facilitating the expansion and maturation of the autophagosome [49]. In this study, we observed that all four environmental stresses could hinder autophagy (Figure 6 C). Specifically, the number of autophagosomes significantly increased under acetic acid and H2O2 conditions. Conversely, the green fluorescent protein (GFP) signal dispersed from the autophagosome or vacuole to the cytoplasm under high temperature and ethanol conditions (Figure 6 C).

### Apoptosis related yeast protein

Yca1p is a metacaspase, similar to caspases found in animals, that is involved in yeast apoptosis, or programmed cell death. It is specifically activated in response to various stress conditions, such as oxidative stress, and helps break down cellular components in a controlled manner [50]. Aif1p (Apoptosis-Inducing Factor 1) is a protein found in Saccharomyces cerevisiae that plays a crucial role in cell death processes. It shares similarities with the mammalian apoptosis-inducing factor (AIF) and is responsible for the yeast’s response to oxidative stress [51]. Mmi1p, the yeast counterpart of mammalian TCTP, is reported to translocate from the cytoplasm to the mitochondria under oxidative stress conditions and participate in the stress-induced cell death pathway [52]. Yca1p, Aif1p, and Mmi1p have been reported to play important roles in various stress conditions. Therefore, we aimed to systematically examine their localization under a common environmental stress condition. In this study, we observed that the cytoplasmic signal of Yca1-2GFP aggregated under high temperature and ethanol-induced conditions. However, we found that the aggregated puncta did not colocalize with mitochondria (Figure 7 A). Interestingly, the cytoplasmic signal of Yca1-2GFP remained unchanged under acetic acid and H2O2-induced conditions (Figure 7 A). Similarly, Aif1-2GFP exhibited similar behavior to Yca1-2GFP under high temperature, ethanol, acetic acid, and H2O2-induced conditions (Figure 7 B). Furthermore, we observed that 2GFP-Mmi1 also aggregated under high temperature and ethanol-induced conditions. However, the aggregated puncta of Mmi1 partially localized to the nucleus under high temperature condition, while the ethanol-induced aggregated puncta of Mmi1 did not localize to the nucleus (Figure 7 C).

**Fig 7.**
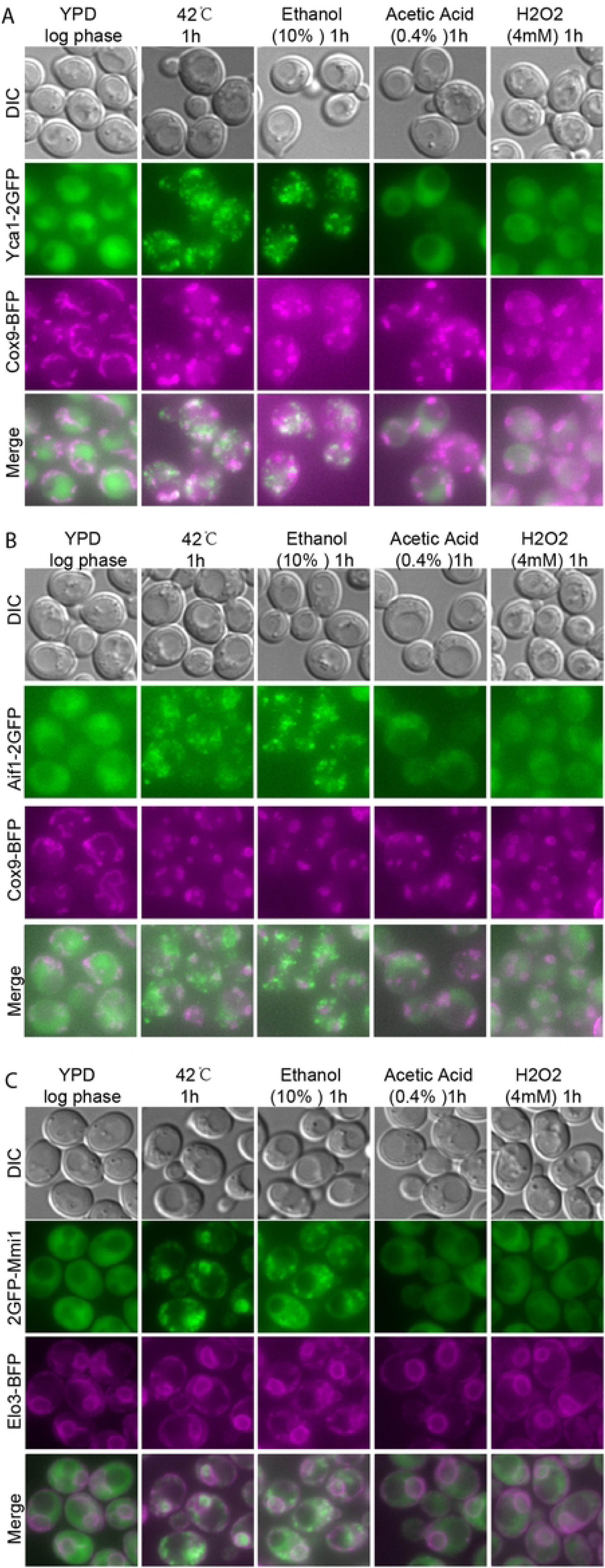
Stress response of apoptosis related-proteins. Images were captured and presented as in Fig 1 (A-C). Colocalization between the Yca1-2GFP and Cox9-mTagBFP (A). Colocalization between the Aif1-2GFP and Cox9-mTagBFP (B). Colocalization between the 2GFP-Mmi1 and Elo3-mTagBFP (C).

## Discussion

Eukaryotic cells experience stress during different stages of their growth and reproductive cycles. To counteract the detrimental effects of various stress conditions, they have evolved a range of strategies. For example, yeast cells undergo multiple stresses, such as thermal, oxidative, ethanol, acetic acid, and/or starvation during biomass propagation. Among these stresses, high temperature is particularly significant during fermentation. When the temperature exceeds 36℃, yeast cells activate a protective response known as the high temperature response (HSR). The increase in temperature can cause proteins to unfold, tangle, and form aggregates, which can harm cell morphology and phenotype. Additionally, stress conditions can lead to damage to the cytoskeleton, resulting in incorrect organelle localization and breakdown of intracellular transport processes [53]. Studies have shown that stress conditions can cause fragmentation of the Golgi system and the endoplasmic reticulum (ER), as well as a decrease in the number of mitochondria and lysosomes [54, 55]. In addition to high temperature, the accumulation of reactive oxygen species (ROS) can cause oxidative damage to nucleic acids, proteins, and lipids, leading to apoptosis and necrosis. ROS can originate from various exogenous sources, such as exposure to hydrogen peroxide (H2O2), superoxide anion, heavy metals, treatment with specific anticancer drugs, or elevated temperature [56]. Another common stressor for yeast cells during fermentation is an elevated concentration of ethanol. Yeast cells have developed mechanisms to effectively deal with the damage caused by increased ethanol levels [56]. Acetic acid, a metabolic byproduct of yeast produced during fermentation, can hinder yeast fermentation and cause widespread cell death when it accumulates excessively [14]. The biochemical markers of cell death induced by acetic acid (20-80mM) were initially characterized as chromatin condensation, phosphatidylserine (PS) exposure, DNA strand breaks, cytochrome c release, and accumulation of reactive oxygen species (ROS) [59, 60]. Moreover, studies have shown that acetic acid-induced apoptosis leads to the fragmentation and deformation of the normal tubular mitochondrial network, resulting in a punctate pattern [61–65]. Apart from mitochondria, the vacuole of yeast also plays a crucial role in acetic acid-induced regulated cell death (AA-RCD), involving vacuolar membrane permeabilization and subsequent release of Pep4p [66]. However, no research has been conducted to examine the structural changes of all membrane-based cell organelles under stress-induced conditions. Therefore, in this study, we systematically examined the structural changes of membrane-based cell organelles under various conditions, including high temperature, oxidative stress, ethanol, and acetic acid-induced conditions. The results of our study are summarized below.

### Mitochondria

In this study, we used various fluorescent protein-based markers and a dye called MitoTracker to label specific structures within the mitochondria. The markers we used were Om14-2GFP, Abf2-2GFP, Afg3-2GFP, Cit1-2GFP, Cox9-mTagBFP, Ndi1-2GFP, and Nuc1-2GFP.

Our findings indicated that under the four different stress conditions mentioned above, the mitochondria exhibited the same structural change, transitioning from a filamentous form to punctate structures. Interestingly, despite this fragmentation, two well-known apoptosis-related proteins, Ndi1p and Nuc1p, remained within the mitochondria. This is in contrast to previous studies that reported Nuc1p translocating from the mitochondria to the nucleus and Ndi1p relocating to the cytoplasm during apoptosis induction [8, 9]. The discrepancy in the observed phenomenon may be attributed to variations in the conditions used to induce apoptosis between our study and previous research.

### ER

In this study, we used three markers, Emc1-2GFP, Elo3-mTagBFP, and GFP-HDEL, to label the endoplasmic reticulum (ER). Our results demonstrate that the signal of the ER resident proteins Emc1p and Elo3p remained unchanged under all four stress conditions mentioned. The HDEL sequence, composed of Histidine-Aspartic acid-Glutamic acid-Leucine, is a specific amino acid sequence found at the C-terminus of certain proteins that are destined for the endoplasmic reticulum (ER). This sequence acts as a signal, ensuring that these proteins are retained within the ER and not secreted or transported elsewhere. Receptors in the Golgi apparatus recognize proteins carrying the HDEL sequence, facilitating their retrieval back to the ER if they escape, thus maintaining their localization within the ER [30]. This system is crucial for the proper functioning of the ER, where these proteins are typically involved in protein folding, processing, and quality control. However, our observations reveal that more than half of the GFP-HDEL signal dispersed from the ER into the cytosol under all four stress conditions. This finding suggests a disruption either in the functionality of the ER or in the communication between the ER and Golgi.

### ER Unfolded Protein Response

The Unfolded Protein Response (UPR) is crucial for survival in situations that result in protein misfolding, such as high temperature, oxidative stress, or mutations in proteins that impede proper folding. A failure in the UPR can lead to cell death due to the toxic accumulation of unfolded proteins. In yeast, Ire1p acts as the primary detector of misfolded proteins in the ER [31]. When unfolded proteins accumulate, Ire1p forms oligomers and undergoes autophosphorylation, activating its endoribonuclease activity. Once activated, Ire1p splices the mRNA of the transcription factor Hac1p. [32]. In this study, we investigated the unfolded protein response of the ER using Ire1-GFP and found that high temperature and oxidative stress can trigger the aggregation of Ire1. However, neither ethanol nor acetic acid could induce the Ire1-mediated ER unfolded protein response. This disparity in stress conditions that elicit the ER unfolded protein response suggests distinct mechanisms underlying the cell stress response.

### Nucleus

In this study, we used several markers, including Nab2-GFP, Nsr1-GFP, Pus1-2GFP, Nhp6a-2GFP, Nma111-2GFP, Ssn8-2GFP, and Mcd1-2GFP, to identify the nucleus. Our findings indicate that during the logarithmic growth phase in YPD medium, all of these resident proteins were localized within the nucleus. However, when exposed to external stresses, they either partially or completely relocated to the cytosol. This phenomenon suggests that while the structure of the nucleus remains relatively unchanged, the behavior of its resident proteins undergoes alterations.

### Vacuole

In this study, we used Pho8-GFP, Ybh3-2GFP, Prc1-2GFP, and Pep4-2GFP to label the vacuole. Pho8 is a vacuolar (lysosomal) alkaline phosphatase [40]. Ybh3p is a yeast protein that contains a BCL-2 homology 3 (BH3) domain, which is a characteristic feature of pro-apoptotic members of the BCL-2 family of proteins. In mammals, BH3-only proteins such as Bid, Bad, and Bim are pro-apoptotic and can initiate the apoptotic cascade by binding to and neutralizing anti-apoptotic BCL-2 family members, or by directly activating pro-apoptotic effectors like Bax and Bak [67]. Previous research has shown that when treated with acetic acid, BXI1/Ybh3 translocates from predominantly vacuolar sites to mitochondria [19] Prc1p is a vacuolar carboxypeptidase Y [41]. Pep4p is cathepsin D (CatD), also known as proteinase A in yeast, and it has been reported to translocate from the vacuole to the cytosol in response to apoptosis induced by H2O2 or acetic acid [14–16]. In our study, during the logarithmic growth phase in YPD medium, all four vacuole markers were localized within the vacuole, with Pho8 showing a higher signal at the vacuole membrane compared to the others. However, under acetic acid-induced conditions, Ybh3-2GFP moved from the inner vacuole to the vacuole membrane, Prc1-2GFP and Pep4-2GFP transferred from the inner vacuole to several unknown puncta and the nuclear ER. Interestingly, high temperature, ethanol, and oxidative stress did not change the localization of any of the four vacuole markers, even though the vacuoles fused to form larger ones under these stress conditions. This phenomenon suggests that during stress conditions, vacuoles fuse together to form larger organelles that may be more efficient in coping with stress. However, it should be noted that the apoptosis-related proteins Ybh3p and Pep4p did not transfer to mitochondria under the four stress conditions we tested.

### Golgi and Endosome

In yeast, the Golgi apparatus and endosomes are both integral components of the cellular trafficking and sorting machinery. They play essential roles in the processing, modification, and transport of proteins and lipids. Although they are distinct organelles, they closely interact to manage the flow of materials within the cell. The Golgi apparatus in yeast cells is an essential organelle involved in the modification, sorting, and packaging of proteins and lipids, similar to more complex eukaryotic cells. However, unlike the single, large Golgi apparatus found in mammalian cells, the yeast Golgi is composed of multiple small, dispersed cisternae throughout the cytoplasm. These individual cisternae are known as early- and late-Golgi [20]. Endosomes are membrane-bound organelles that serve as sorting stations within the cell. They direct incoming material from the plasma membrane or Golgi to various destinations, such as the vacuole. The Golgi and endosomes are interconnected through vesicular trafficking. In this study, we used specific labels to visualize different compartments: Vrg4-2GFP and Sys1-2GFP for the early-Golgi, Chs5-GFP and Sec7-2GFP for the late-Golgi, and Snf7-2GFP and Vps4-2GFP for the late-endosome. We observed that under high temperature and ethanol-induced conditions, the puncta signal of Vrg4-2GFP and Sys1-2GFP in the early-Golgi faded away. Additionally, the number of puncta for these proteins decreased under acetic acid and H2O2-induced conditions. Regarding the late-Golgi, we found that the puncta of Chs5 and Sec7 completely vanished under H2O2-induced conditions. However, under high temperature conditions, the GFP puncta clustered together, and under ethanol-induced conditions, the GFP signal significantly faded away. Furthermore, under acetic acid-induced conditions, the GFP puncta became larger and lighter, especially in the case of Sec7-2GFP. These results indicate that the structural changes in the Golgi during stress conditions may depend on the specific stressors. In relation to the late endosome, Snf7-GFP formed aggregated puncta during the logarithmic growth phase in YPD medium, and then translocated to the vacuole membrane under all four stress conditions mentioned above. On the other hand, Vps4-GFP appeared as small puncta during the logarithmic growth phase in YPD medium, but disappeared under high temperature, ethanol, and acetic acid-induced conditions. The signal of Vps4-GFP did not change significantly under oxidative stress. These findings suggest that the structural changes in the late endosome may not be consistent among specific stressors, and different proteins may undergo varying changes.

### Lipid droplet, Peroxisome, and Autophagosome

The synthesis of lipids in cells can be affected by stress conditions, particularly in yeast cells. Our study found that high temperature can inhibit lipid synthesis, while ethanol, acetic acid, and oxidative stress induced by H2O2 can stimulate lipid synthesis. Of these stressors, acetic acid and oxidative stress were especially effective. This suggests that stress conditions may lead to lipid peroxidation and the degradation of membrane lipids. To counteract this, cells may increase the production of specific lipids to repair and maintain membrane integrity. Peroxisomes are dynamic organelles involved in lipid metabolism, detoxification of reactive oxygen species (ROS), and the breakdown of fatty acids. Under stress conditions, peroxisomes play a crucial role in maintaining cellular homeostasis and responding to environmental changes. Our study observed that high temperature and ethanol caused the aggregation of Pex1-2GFP, indicating an effect on peroxisomes. However, acetic acid and H2O2 had little impact on peroxisomes. Autophagy is a cellular process that involves the degradation and recycling of damaged or unnecessary cellular components. It is critical for maintaining cellular homeostasis, particularly under stress conditions. When cells are exposed to stressors, autophagy is often upregulated as a survival mechanism to help the cell cope and prevent damage. When essential nutrients such as amino acids, glucose, or lipids are deprived, autophagy is activated to form autophagosomes. These autophagosomes break down cellular components and recycle them to generate energy and building blocks for the cell. In our study, we observed that after 2 hours of nitrogen starvation, most of the autophagosomes translocated into vacuoles. However, under high temperature, ethanol, acetic acid, and oxidative conditions, the autophagy pathway was blocked. This indicates that while autophagy is essential for cells to deal with stress, severe stress can disrupt this pathway.

## Acknowledgments

We are grateful to Xie Zhi-ping (Shanghai Jiao Tong University) for providing By4741 strain and fluorescent plasmids.

